# Genetic diversity and demographic history of the largest remaining migratory population of blue wildebeest (*Connochaetes taurinus taurinus*) in southern Africa

**DOI:** 10.1101/2024.09.05.611351

**Authors:** Stephanie J. Szarmach, Katherine C. Teeter, Jassiel M’soka, Egil Dröge, Hellen Ndakala, Clive Chifunte, Matthew S. Becker, Alec R. Lindsay

## Abstract

The blue wildebeest (*Connochaetes taurinus taurinus*) is a keystone species in the savannahs of southern Africa, where it maintains shortgrass plains and serves as an important prey source for large carnivores. Despite being the second largest migratory wildebeest population, the wildebeest of the Greater Liuwa Ecosystem (GLE) of western Zambia have remained largely unstudied, until recently. While studies have increased understanding of recent demography, migration, and population limiting factors, the level of genetic diversity, patterns of gene flow, and long-term demographic history of blue wildebeest in the GLE remains unknown. Most genetic studies of wildebeest have focused on small, heavily-managed populations, rather than large, migratory populations of high conservation significance. We used restriction-site associated DNA sequencing (RAD-seq) to assess genetic diversity, population structure, and demographic history of blue wildebeest in the GLE. Using SNPs from 1,730 loci genotyped across 75 individuals, we found moderate levels of genetic diversity in GLE blue wildebeest (*H*e = 0.210), no evidence of inbreeding (*F*IS = 0.033), and an effective population size of about one tenth the estimated population size. No genetic population structure was evident within the GLE. Analyses of the site frequency spectrum found signatures of expansion during the Middle Pleistocene followed by population decline in the Late Pleistocene and early Holocene, a pattern previously observed in other African ungulates. These results will supplement field studies in developing effective conservation plans for wildebeest as they face continued and increasing threats of habitat loss, poaching, and other human impacts across their remaining range.

## INTRODUCTION

The blue wildebeest (*Connochaetes taurinus*) is a large antelope that acts as a keystone species in the savannahs of southern and eastern Africa. Many wildebeest populations migrate seasonally in response to precipitation, availability of high-quality forage, and predation risk [1–4]. Grazing by wildebeest maintains shortgrass plains and facilitates nutrient cycling [5–7], which in turn supports communities of other grazing mammals, insects, and birds that depend on small herbaceous plants [8]. Carnivores, such as lions (*Panthera leo*), hyenas (*Crocuta crocuta*), wild dogs (*Lycaon pictus*), and cheetahs (*Acinonyx jubatus*), often depend on wildebeest as an important prey source [9–12]. Because of the significant ecological roles of wildebeest, changes in migration dynamics or population size could have dramatic impacts on other species and communities in the African grasslands [13].

In many parts of the wildebeest’s range, habitat destruction and fragmentation, obstruction of migration routes, and poaching have impacted wildebeest population size and movement [14]. The conversion of grassland to agricultural fields has reduced available foraging area, contributing to population declines [15,16] and the construction of fences and roads has blocked migration routes and caused previously migratory populations to become sedentary [17,18]. Poaching for bushmeat, driven by deteriorating economic conditions in many communities in and around protected area networks, also threatens wildebeest populations [19–21]. To better understand and address the impact of these threats, there is a need to detect and evaluate their effects on populations.

In addition to experiencing recent declines, wildebeest populations may have undergone fluctuations in the more distant past due to changes in climate or human activity. Genetic analysis of other African ungulates, including African buffalo (*Syncerus caffer*) [22–24], African elephant (*Loxodonta africana*) [25], and hippopotamus (*Hippopotamus amphibius*) [26,27], revealed population declines during the late Pleistocene and early Holocene. These declines coincide with increases in human population size and periods of aridity on the African continent, during which longer dry seasons led to drought and changes in water levels and vegetation types [28–30]. Because many wildebeest populations migrate in response to rainfall and vegetation growth and populations have declined during recent droughts, wildebeest likely experienced similar declines during these two epochs [4,19].

Large population declines during the Late Pleistocene and Holocene may have affected the level of genetic diversity found in the species today, and an understanding of these changes could help in evaluating future demographic effects of climate change. Today, the dryland ecosystems comprising most of the remaining wildebeest range are characterized by extreme seasonality in resource abundance and distribution, making mobility and access to these resources key, particularly in the face of climate change [31]. Understanding how continued habitat fragmentation, loss of migratory routes, and population declines affect genetic diversity and the ability of wildebeest to adapt to the rapidly changing environments of the Anthropocene is therefore of key importance.

The Greater Liuwa Ecosystem (GLE), located in western Zambia and encompassing Liuwa Plain National Park (3,660 km^2^) and the Upper West Zambezi Game Management Area (22,156 km^2^; Figure 1), supports a large, migratory blue wildebeest population. This flat, open grassland experiences extensive seasonal flooding, which drives the wildebeest to migrate between the drier, southern portion of the GLE, where they spend the wet season (November – April), and the previously flooded northern areas for the dry season (May – October) [32,33]. Though this is thought to be the second largest wildebeest migration in Africa, little was known about this population until recently, after it experienced a significant decline. Poaching, driven by regional politics and civil war in neighboring Angola, led to severe declines of many species in the GLE, including wildebeest, and notably led to the extirpation of lions [32,34].

**Fig. 1.**
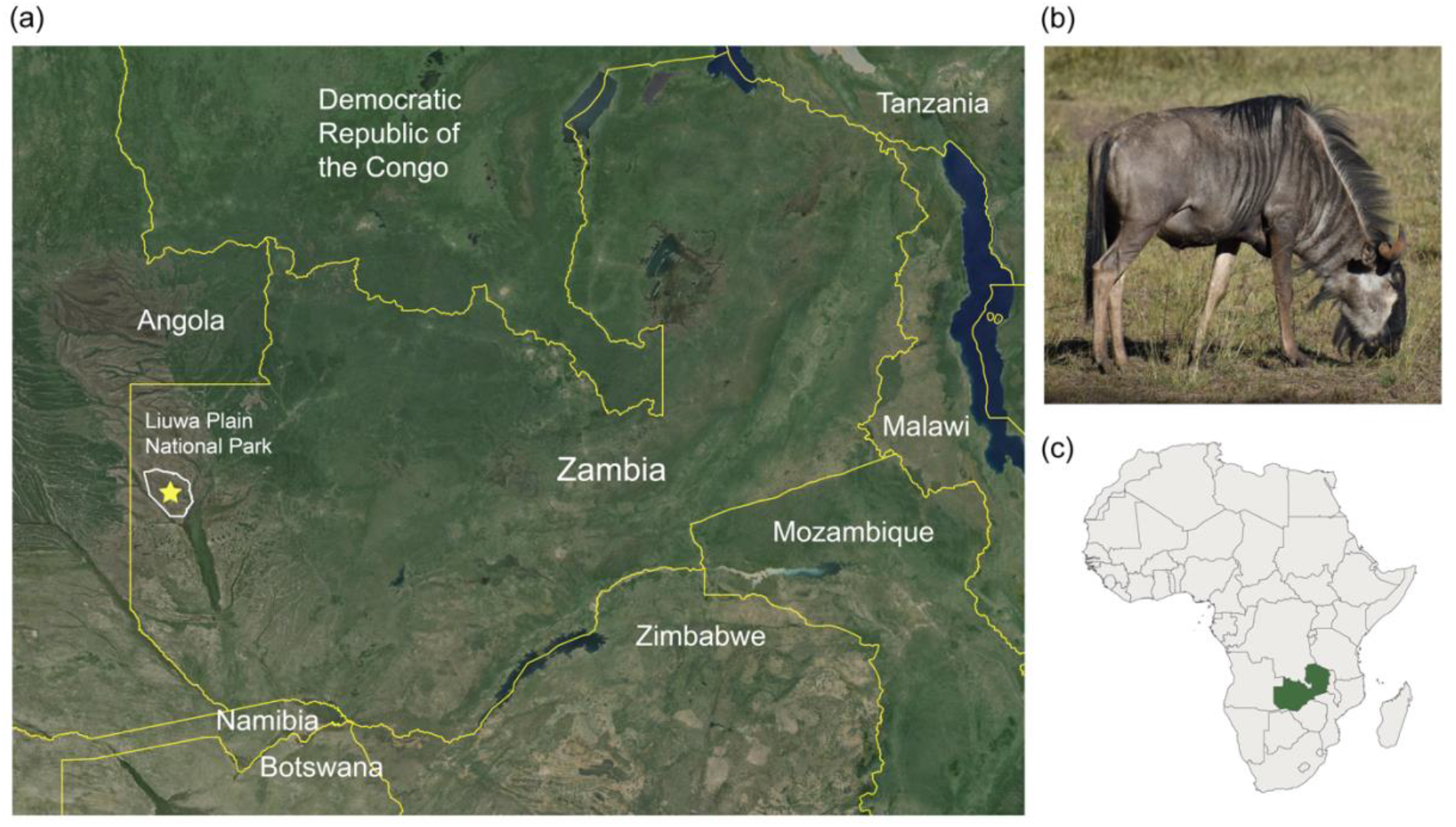
Location of study area. (a) Satellite map of Zambia and surrounding area with Liuwa Plain National Park, where blue wildebeest tissue samples were collected for this study, outlined in white and marked with a star. Basemap: Google Satellite (obtained through QuickMapServices QGIS plugin), Map data © 2024 Google. (b) Image of a blue wildebeest (Photograph by S. Szarmach). (c) Location of Zambia (green) within Africa.

Additionally, while the wildebeest are thought to have historically migrated between Zambia and Angola, since the Angolan civil war (1975-2002) no transboundary migration has been observed and radio-tracking has shown wildebeest movements to be largely confined to the GLE [33].

Restoration efforts in Liuwa began in 2003 with the signing of a 20-year management agreement between the government of Zambia and African Parks Network. Ecosystem recovery efforts have resulted in growth and stabilization of the wildebeest population, from an estimated 15,000 individuals in 2001 to 46,000 in 2013 [33–35]. Studies of movement, demography, environmental influences, and predator-prey dynamics [11,32,33,36] have improved understanding of wildebeest population dynamics in the GLE, but the amount of genetic diversity present in the population and its long-term demographic history remain unknown. Past changes in population size have important implications for conservation in the present day, because historical bottlenecks can reduce genetic diversity and limit future adaptive potential, and the comparison of present day to past effective population sizes can improve understanding of current conservation risks for the population [37,38].

Despite the wildebeest being an iconic species of the African savannahs with high conservation significance, until very recently [39], few genetic studies of wildebeest had been published. Studies utilizing molecular markers can inform wildlife management by delineating conservation units, revealing inbreeding, and divulging historical patterns of movement and demographic change [40,41]. Most previous genetic studies of wildebeest focused on heavily managed populations in South Africa, and few evaluated the potential impacts of habitat loss, fragmentation, and population declines occurring across remaining wildebeest range. Early studies assessing wildebeest genetic diversity used allozymes, microsatellites or mitochondrial DNA primarily to inform management of small populations of wildebeest on game ranches in South Africa [42–44]. Microsatellite and mtDNA studies that included samples from a greater part of the blue wildebeest’s range identified high levels of genetic divergence between wildebeest in southern versus eastern Africa, as well as lower levels of significant differentiation between pairs of populations within these regions [45,46].

More recently, whole-genome resequencing of blue wildebeest from much of the species’ range found significant genetic differentiation between all subspecies of blue wildebeest, as well as between local populations of the western white bearded (*C. t. mearnsi*) subspecies in eastern Africa and the brindled (*C. t. taurinus*) subspecies in southern Africa [39]. Importantly, Liu et al. [39] found that wildebeest populations where migration had recently been disrupted showed lower genetic diversity and higher levels of inbreeding compared to fully migratory populations. No previous genetic studies have included wildebeest from the GLE, despite this being one of the largest migratory wildebeest populations. The effects of past changes in population size, both historical and recent, on genetic diversity in this recovering migratory population remain unknown.

We assessed genetic diversity, population structure, and demographic history of the GLE blue wildebeest population using a genome-wide SNP dataset generated through restriction-site associated DNA sequencing (RAD-seq). We first evaluated the current level of genetic diversity present in the GLE wildebeest population, estimated contemporary effective population size, and assessed whether genetic population structure is present. We used both model-based (STRUCTURE) and model-free (PCA) approaches to determine whether multiple populations exhibiting significant differences in SNP allele frequencies are present within the GLE, or whether this region is occupied by a single, panmictic population. We then inferred the past demographic history of the GLE blue wildebeest population and tested the hypothesis that genetic signatures of population contraction are present due to past climate change and anthropogenic pressures. To do this, we tested demographic models using the coalescent simulator fastsimcoal2 [47] and implemented the model-independent stairway plot method [48,49] to estimate effective population size over time based on the site frequency spectrum.

## MATERIALS AND METHODS

### Sample collection and DNA extraction

As part of long-term field studies conducted with the Zambia Department of National Parks and Wildlife (DNPW) and African Parks, we collected 97 tissue samples in the GLE between 2011−2014 from wildebeest that were anesthetized for radio collaring as part of long-term studies, or from opportunistically sampled wildebeest carcasses detected during the course of fieldwork. These tissue samples were transferred to ethanol for storage. All animal handling procedures were performed following animal welfare standards and protocols required by the Zambia Department of Veterinary and Livestock Services and the DNPW and were approved by the Montana State University IACUC. Samples were exported with Zambia Wildlife Authority Permit #7830 and imported with USDA Veterinary Permit #109174. We extracted genomic DNA from the tissues using the DNeasy Blood & Tissue Kit (Qiagen, Valencia, CA, USA) and quantified the concentration and purity of the extracted DNA using a Nanodrop 2000c. DNA extractions with a concentration <10 ng/μL were combined with DNA from a second extraction from the same tissue sample, if available, and concentrated using vacuum centrifugation.

### RAD library preparation and sequencing

We prepared double-digest restriction site associated DNA (ddRAD) libraries following the protocol of DaCosta and Sorenson [50]. First, ∼1 µg of genomic DNA from each wildebeest was digested using the restriction enzymes *EcoR*I-HF and *Sbf*I-HF (New England Biolabs, Ipswich, MA, USA). Unique combinations of barcoded adapters were ligated to the digested DNA from each individual, so that samples could later be pooled for sequencing. Restriction fragments within a size range of 200−400 bp were selected through gel excision, and DNA was purified using a Qiagen MinElute Gel Extraction Kit. The size-selected DNA fragments were PCR amplified using Phusion High-Fidelity PCR Master Mix (Thermo Scientific, Waltham, MA, USA). PCR products were purified with AMPure XP SPRI beads (Beckman Coulter, Brea, CA, USA). The concentrations of the purified PCR products were quantified using real-time PCR, and samples were pooled in equimolar amounts, with the exception of some low-concentration samples. Ultimately, 83 samples were included in the multiplexed library, which was sequenced on one lane of an Illumina HiSeq 2500 Sequencing System (Tufts University, Medford, MA, USA), generating single-end reads of 100 bp in length.

### Alignment, genotyping, and quality filtering

After sequencing, we used FastQC version 0.11.7 to assess per-base sequence quality for the reads [51]. We used a computational pipeline developed by DaCosta and Sorenson [50] for analyzing ddRAD-seq data to process the raw sequence reads (Python scripts available at https://github.com/BU-RAD-seq/ddRAD-seq-Pipeline). After correcting sequencing errors in restriction sites and barcodes, reads were assigned to individual wildebeest based on the combination of barcodes associated with the read. The demultiplexed sequence data are available from the NCBI Sequence Read Archive (SRA) under the following accession numbers: BioProject ID PRJNA588629; BioSample IDs SAMN13254976−SAMN13255074. After demultiplexing, identical sequences were collapsed and reads with low average Phred quality scores (Q < 20) were filtered out. The condensed and filtered reads were then clustered into putative loci across individuals based on a similarity threshold of 0.85 using the UCLUST algorithm in the program USEARCH version 5 [52]. A representative sequence from each cluster was aligned to a reference genome using the blastn program in BLAST version 2.8.0 [53] and the following settings: evalue = 0.0001, word_size = 11, gapopen = 5, gapextend = 2, penalty = -3, reward = 1, and dust = yes). Sequences were aligned to two reference genomes: version 4.0 of the domestic sheep (*Ovis aries*) genome (2015; Assembly GCA_000298735.2), downloaded from NCBI, and the blue wildebeest genome sequenced and assembled by Chen et al. [54], downloaded from http://animal.nwsuaf.edu.cn/code/index.php/Ruminantia. Separate clusters that generated BLAST hits to the same genomic location were combined into a single cluster, and those that did not generate hits to the reference genome were retained but marked as anonymous loci. Reads were aligned within each cluster using MUSCLE version 3.8.31 [55].

We genotyped individuals at the identified loci using DaCosta and Sorenson’s RADGenotypes.py script [50], which calls individuals as homozygous for a SNP if >93% of reads exhibit the same allele, and as heterozygous if >29% of reads show a second allele. When a lower proportion of reads exhibited a minor allele, the genotype was flagged as a “provisional heterozygote” (20-29% of reads) or “ambiguous” (7-20%) and was required to pass additional examination and quality filtering before being included in the final set of SNPs. A custom R script utilizing the tidyverse collection of packages [56,57] was used to remove loci with substantial amounts of missing data and putative “paralogous loci” formed by erroneous clustering of sequences originating from repetitive elements in the genome (R script available at https://github.com/sszarmach/NMU-wildebeest). Samples with an average sequencing depth of <10 reads per locus and loci with <5 reads in >10% of samples were removed, since low sequencing depth can lead to genotyping errors [58]. We filtered out loci that significantly departed from Hardy-Weinberg equilibrium based on a chi-square goodness-of-fit test (α = 0.05), because such deviation can result from null alleles (heterozygote deficiency) or incorrectly clustered paralogous loci (heterozygote excess). To eliminate putative paralogous loci, we also removed clusters that exhibited more than two alleles, that had abnormally high read depths (mean depth >700), that had >4 SNPs, or that generated multiple high-quality BLAST hits to the reference genome [59,60]. For all downstream analyses except inference of demographic history using the site frequency spectrum, a minor allele frequency filter (MAF >0.01) was applied to remove loci containing singleton SNPs, which could be an artifact of sequencing error. These quality filtering steps are outlined in Supplementary Table 1.

### Genetic diversity

A final data set of 1,730 polymorphic loci genotyped across 75 individuals was used to calculate summary statistics describing genetic diversity in GLE blue wildebeest. We randomly drew one SNP from each RAD locus to avoid the inclusion of multiple SNPs that were linked due to chromosomal proximity. We used PGDSpider version 2.1.1.5 to convert the SNP data file into the correct format for each analysis program [61]. Allele frequencies and expected and observed heterozygosities were calculated using the R package adegenet [62,63]. We tested SNPs for deviation from Hardy-Weinberg equilibrium using the hw.test function in the R package pegas (Fisher’s exact test with 1,000 MCMC replicates) [64]. *P*-values from the Hardy-Weinberg tests were corrected for multiple comparisons using the Benjamini and Hochberg method in the R package sgof [65–67]. The inbreeding coefficient, *FIS*, was estimated using the R package hierfstat [68].

### Estimation of effective population size

We estimated contemporary effective population size (*Ne*) for GLE blue wildebeest using the bias-corrected linkage disequilibrium method [69,70] implemented in NeEstimator version 2.1 [71] assuming a random-mating model. 95% confidence intervals for *Ne* were constructed using the parametric method of Waples [69] and the jackknifing method of Jones et al. [72], which accounts for pseudoreplication resulting from physically linked loci and is recommended for data sets including large numbers of SNPs.

### Population structure

We explored whether genetic structure is present within the GLE wildebeest population using the model-based clustering approach implemented in STRUCTURE version 2.3.4 [73–76]. We assumed the admixture model and correlated allele frequencies [74,77]. Ten independent runs were performed for each number of genetic clusters tested (*K* = 1 to *K* = 4), and each run contained 10,000 burn-in steps and 100,000 MCMC steps. We used STRUCTURE HARVESTER web version 0.6.94 [78,79] to summarize STRUCTURE results, and the best *K* value was selected based on the log probability of the data given the number of clusters (ln Pr(X|*K*)) and the Evanno Δ*K* method [80]. Visualizations of the STRUCTURE output were created using the R package and Shiny app pophelper [81]. We also examined genetic structure using a non-model-based approach by performing a principal components analysis (PCA) in the R package adegenet [62].

### Demographic history

Using the coalescent simulator fastsimcoal version 2.6 [47] we evaluated five models of historical demographic change: constant population size, expansion, decline, bottleneck, and expansion followed by decline (hereafter “expansion-decline”; Figure 2). Fastsimcoal2 simulates the expected site frequency spectrum (SFS) for a demographic model provided by the user and computes the composite likelihood of the SFS observed from the data given this model. We computed the folded site frequency spectrum from the minor allele frequencies of all sites in 1,831 loci (2,946 SNPs and 172,603 monomorphic sites) that passed quality filtering without applying the minor allele frequency filter, as excluding singletons can bias inferences drawn from the site frequency spectrum [82].

**Fig. 2.**
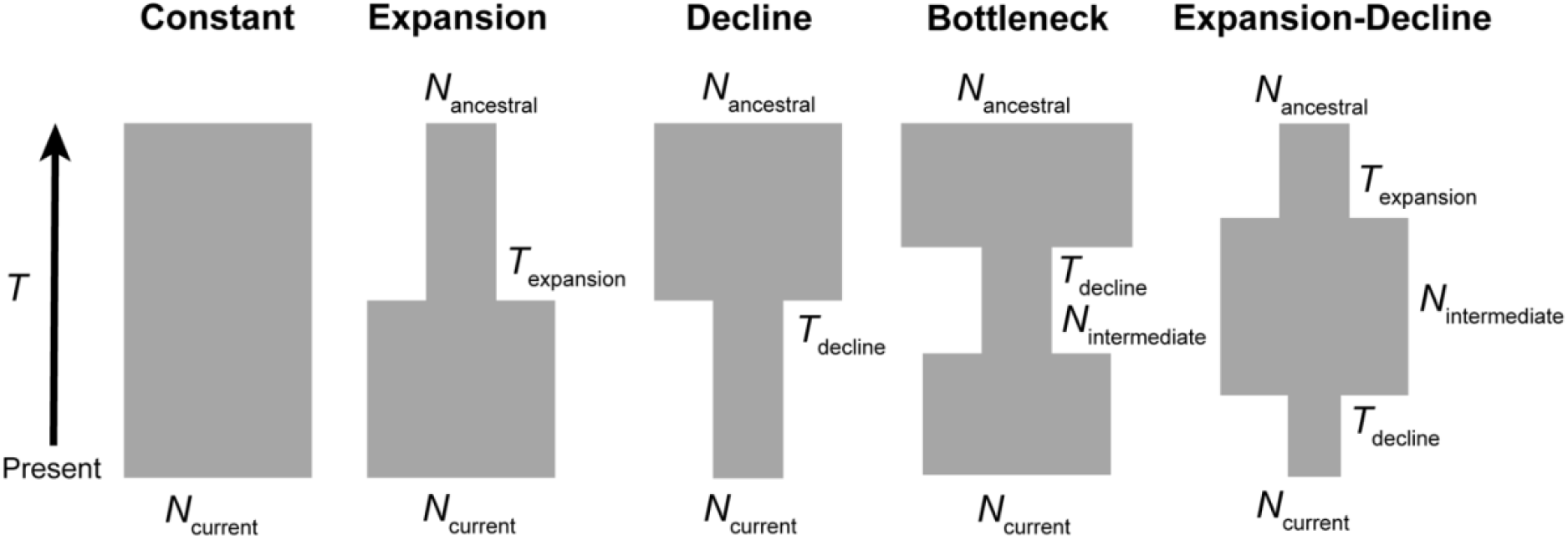
Schematic of the five demographic scenarios modeled in the coalescent simulator fastsimcoal2. For each model, parameters describing effective population size (*N*) and select times before present (*T*) were estimated.

For each model, we ran 50 replicate simulations, each with randomly selected initial parameter values, to increase the likelihood that the global maximum solution was discovered. Each run included 100,000 coalescent simulations and 40 cycles of Brent’s ECM algorithm, which is used to select parameters for the model that optimize the composite likelihood [47]. We provided broad parameter search ranges of 10–100,000 gene copies (haploid *N*) for all *N*e estimates, except *N*ancestral in the decline and bottleneck models and *N*mid in the expansion-decline model, which were set to 10–1,000,000 gene copies. The search range for all event time parameters was 10-100,000 generations. The mutation rate was set to 1.7x10^-8^ substitutions per base pair per generation, based on the average mammalian mutation rate (2.22x10^-9^ substitutions per base pair per year) [83] and the generation time of wildebeest (7.5 years) [84]. For each demographic scenario, the replicate with the highest likelihood was retained as the best model. Fastsimcoal2 handles effective population size as the number of gene copies, so *N*e was converted to the number of diploid individuals by dividing all estimated *N*e parameters by two. Time parameters were converted from number of generations to years before present by multiplying *T* estimates by a generation time of 7.5 years. We calculated Akaike Information Criterion (AIC) values [85] for the models with the highest maximum composite likelihood for each demographic scenario and compared the demographic models using Akaike weights (AICw) calculated using the akaike.weights function in the qpcR R package [86].

In addition to the model-based approach implemented in fastsimcoal2, we also inferred historical changes in population size using the model-independent stairway plot method. Stairway Plot 2 uses a flexible multi-epoch model to estimate population size over different time intervals based on the site frequency spectrum [48,49]. The same folded site frequency spectrum, mutation rate, and generation time used in the fastsimcoal2 analysis were used to generate the stairway plot. Confidence intervals were created based on 200 bootstrapped site frequency spectra drawn from the observed SFS.

## RESULTS

Sequencing produced ∼220 million reads that passed initial Illumina quality filtering, and demultiplexing resulted in an average of 2.2 million reads assigned to each wildebeest. After these reads were filtered for quality and clustered by similarity, 22,483 RAD loci were identified. A similar number of loci aligned to each reference genome: 19,216 (85.5%) of sequences generated at least one BLAST hit to the sheep genome, and 19,267 (85.7%) of loci aligned to the blue wildebeest genome. Eight wildebeest samples were removed due to low sequencing depth (<10 reads per locus), leaving 75 wildebeest that were genotyped. There were 3,417 polymorphic loci that were genotyped in >90% of samples. Many putatively paralogous loci were removed from the dataset, likely the result of the large proportion of repetitive sequence in bovid genomes (∼50%) [54,87]. After all quality filtering (Supplementary Table 1), the final dataset included 1,730 high-quality polymorphic RAD loci (mean length = 96 bp, mean depth = 355X) that passed a minor allele frequency (MAF) threshold of 0.01, containing 2,921 SNPs. The number of loci that aligned to each chromosome of the sheep genome was correlated with the length of the chromosome (Spearman’s ρ = 0.683, *P* < 0.001; Supplementary Figure 1), indicating that the dataset contains loci evenly spaced throughout the genome.

We estimated genetic diversity and effective population size for GLE blue wildebeest (*n* = 75) using 1,730 biallelic SNPs, with one SNP randomly drawn from each RAD locus. After correcting for multiple comparisons, no SNPs in the final dataset significantly deviated from Hardy-Weinberg equilibrium (α = 0.05). The minor allele frequency per SNP ranged from 0.01 to 0.50, with a mean of 0.14 (SD = 0.13). The observed heterozygosity of the SNPs ranged from 0 to 0.597, and the average observed heterozygosity of GLE blue wildebeest was 0.204 (SD = 0.150; Figure 3). Expected heterozygosity per SNP ranged from 0.013 to 0.504, and the average expected heterozygosity was 0.209 (SD = 0.151). The inbreeding coefficient (*FIS*) for the population was 0.031 (95% CI: 0.026 – 0.038). The estimated effective population size (*Ne*) was 5,075 (parametric 95% CI = 3,360–10,322; jackknifing 95% CI = 800–∞).

**Fig. 3.**
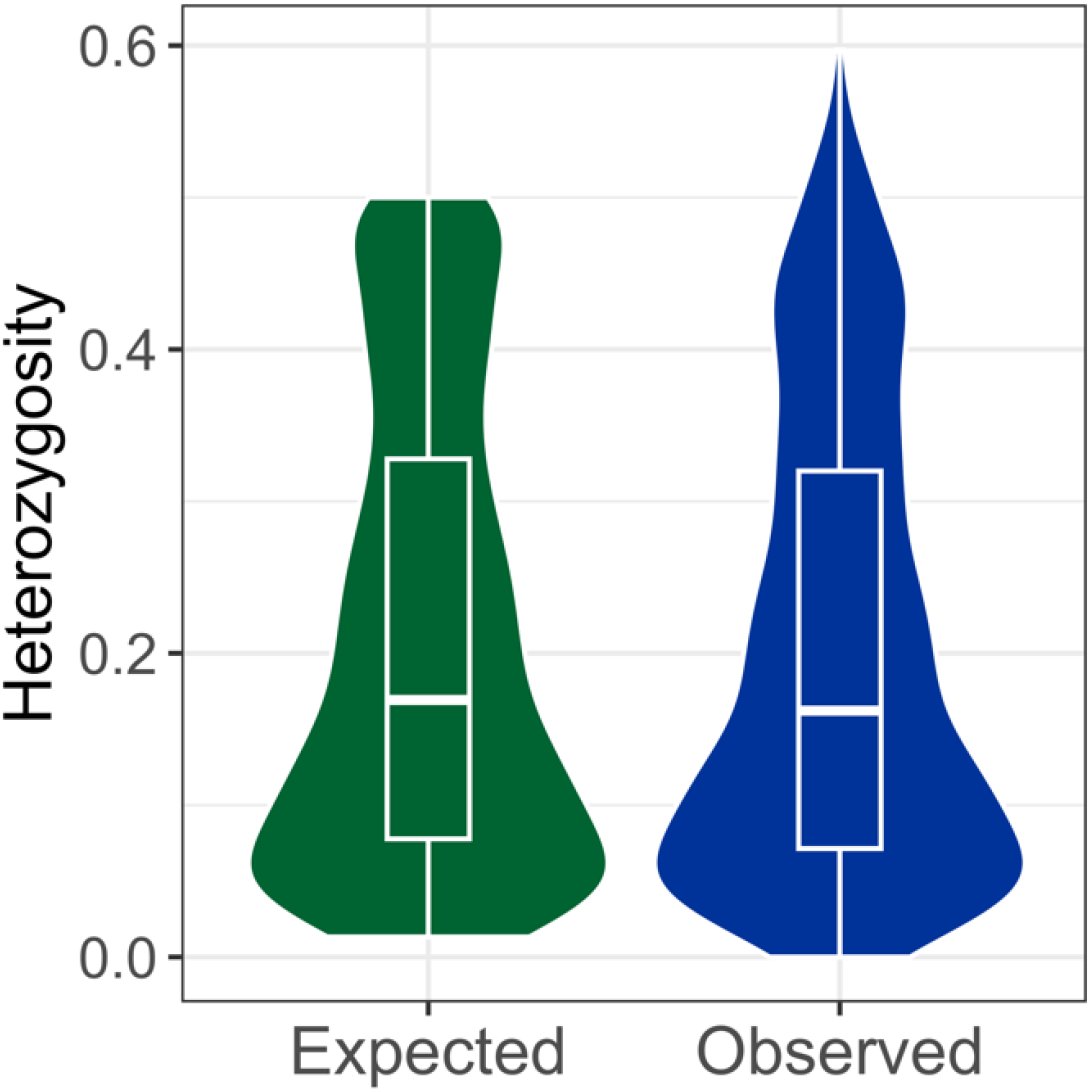
Expected and observed heterozygosity for 1,730 SNPs genotyped in Greater Liuwa Ecosystem blue wildebeest (*n* = 75). Boxplots show median and interquartile range.

We did not detect any genetic population structure within the GLE blue wildebeest. The most likely number of clusters identified using STRUCTURE was *K* = 1 (Supplementary Table 2), and for each *K* value >1 that was tested, almost all individuals were assigned membership to multiple clusters (Figure 4a; Supplementary Figure 2). Principal component analysis also did not show any genetic clustering of subpopulations within the GLE (Figure 4b). Rather, the GLE blue wildebeest seem to compose a single, panmictic population.

**Fig. 4.**
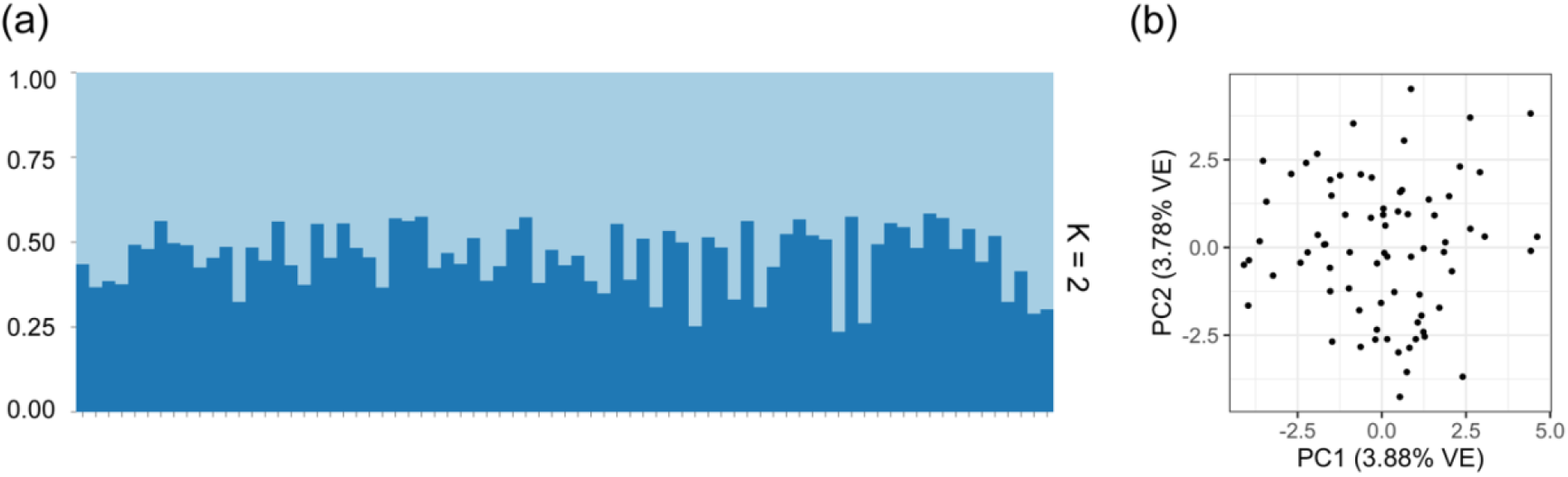
Blue wildebeest show no signs of population subdivision within the Greater Liuwa Ecosystem, based on (a) model-based clustering in STRUCTURE (shown are membership coefficients for *K*=2 clusters for 75 individuals), and (b) principal component analysis (PCA) based on 1,730 SNPs.

The results of both the model-based fastsimcoal2 analysis and model-independent stairway plot analysis suggest that the GLE blue wildebeest experienced population expansion during the Pleistocene followed by large declines throughout the Late Pleistocene and Holocene. Of the five demographic models simulated in fastsimcoal2 (Figure 2), the expansion-decline model had the highest composite likelihood score and Akaike weight (*w*i; Table 1). The parameters of the highest-likelihood expansion-decline model suggest a population increase from *N*e=26,800 to *N*e=92,700 occurring 635,000 years ago, followed by a population decline to *N*e=13,400 occurring 15,480 years ago (Supplementary Table 3). The stairway plot exhibited similar changes in population size as the fastsimcoal2 expansion-decline model (Figure 5), with an increase in population size from *N*e=30,000 to *N*e=90,000 from 1,000,000-500,000 years ago, followed by a sharp decline to *N*e=11,000 from 25,000-3,000 years ago.

**Fig. 5.**
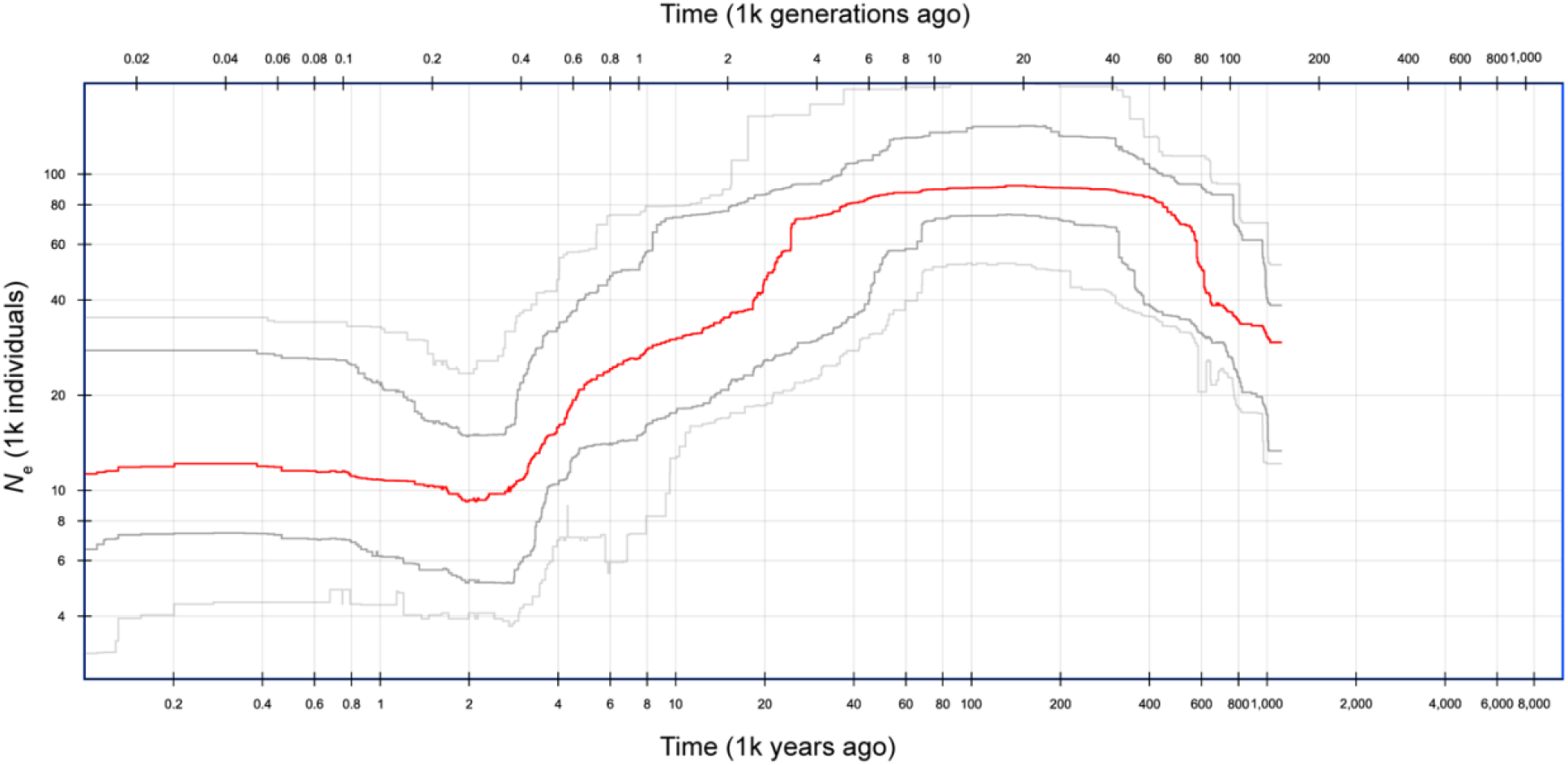
Stairway plot representing inferred changes in effective population size (*N*e) over time for the Greater Liuwa Ecosystem blue wildebeest population. The red line shows the median estimated effective population size, the dark grey lines encompass the 75% CI, and the light gray lines contain the 95% CI, determined from 200 bootstrapped site frequency spectrums.

**Table 1.** Comparison of likelihood scores and Akaike information criterion (AIC) values for the highest likelihood model produced for each demographic scenario tested using the coalescent simulator fastsimcoal2. The best supported model is shown in bold.

| Model | Log likelihood | $k$ | AIC | ΔAIC | Akaike weight ($w_i$) |
| --- | --- | --- | --- | --- | --- |
| Constant size | -11,513.55 | 1 | 23,029.10 | 120.70 | 0.00 |
| Expansion | -11,514.12 | 3 | 23,034.24 | 125.84 | 0.00 |
| Decline | -11,463.62 | 3 | 22,933.25 | 24.85 | 0.00 |
| Bottleneck | -11,462.90 | 4 | 22,933.79 | 25.39 | 0.00 |
| **Expansion-Decline** | **-11,449.20** | **5** | **22,908.40** | **0.00** | **1.00** |

## DISCUSSION

Our results show that a single, panmictic blue wildebeest population occupies the GLE, possessing moderate levels of genetic diversity, no evidence of inbreeding, and no genetic population structure. Though relatively few population genomic studies of wild ungulates have measured SNP-based heterozygosity, the level of heterozygosity seen in GLE blue wildebeest (average per-SNP *Ho* = 0.204) was slightly higher than average per-SNP heterozygosity in another African antelope, the springbok (*Antidorcas marsupialis*; *Ho* = 0.15) [88], and similar to that seen in pronghorn (*Antilocapra americana*; *Ho* = 0.21) [89]. Heterozygosity in GLE wildebeest was slightly lower than that found in African forest elephants (*Loxodonta cyclotis*; *Ho* = 0.27) [90], although the elephant value was averaged over multiple sampling locations. One other study has reported per-SNP heterozygosity for reduced-representation genomic data from blue wildebeest (∼20,000 SNPs generated using DArT-seq) [91]. From their dataset generated for wildebeest from a South African private game ranch, van Deventer et al. [91] found a mean minor allele frequency (0.19) and average observed heterozygosity (0.19) similar to those of the GLE blue wildebeest. However, in the South African population, observed heterozygosity was significantly lower than expected heterozygosity (0.27), potentially a sign of inbreeding that was not seen in the GLE blue wildebeest.

While direct comparison to other previously published studies of genetic variation in blue wildebeest is more challenging, especially those that utilized allozymes, microsatellites, and mitochondrial DNA, the GLE blue wildebeest seem to have relatively greater genetic diversity than some previously studied populations. Researchers found low allozyme polymorphism and low heterozygosity in wildebeest from Tanzania and Kenya [92], and low mtDNA nucleotide diversity [42] and allozyme heterozygosity [43] in South African wildebeest. Many of these early studies focused on smaller, more heavily managed wildebeest populations that likely were experiencing founder effects resulting from translocations of individuals from one reserve to another. In contrast, wildebeest in the GLE are free-ranging and do not have a history of translocations or intensive management. In a recent study of wildebeest populations throughout their range using whole-genome data, Liu et al. [39] found lower genome-wide heterozygosity estimates and more runs of homozygosity (a sign of past inbreeding) in nonmigratory wildebeest populations compared to migratory ones. Though the GLE wildebeest numbers declined drastically in the early 2000’s and the migration no longer extends into Angola, it does not seem that these changes have resulted in low genetic diversity in Liuwa compared to other populations.

The GLE blue wildebeest exhibited no signs of inbreeding. Though *F*IS was significantly greater than zero (0.033; 95% CI: 0.026 – 0.039), indicating a slight excess of homozygosity, this low value does not indicate extensive nonrandom mating in the population. Similar values of the inbreeding coefficient have been seen in springbok (F*IS* = 0.018 – 0.032) from several populations in South Africa [88]. While the GLE blue wildebeest population experienced significant declines in the early 2000’s [34], it seems these declines were not great enough and did not last long enough (occurring over a few generations) to lead to inbreeding and reduction of genetic diversity in the population. We also found no evidence of population structure using either STRUCTURE or PCA, indicating that a single, panmictic wildebeest population inhabits the GLE. Visual observation and GPS tracking of collared wildebeest have found that the GLE wildebeest spend the wet season in a concentrated area in the southern portion of the park [33], and our genetic results corroborate that there is likely no spatial segregation of individuals during breeding.

Our estimate of the current effective population size (*N*e), the size of an idealized population that would experience the same amount of genetic drift as the population being studied [93], was 5,075 (95% CI = 3,360–10,322). Estimating a population’s effective size is important for conservation, because a population with low *N*e could lose genetic diversity faster than expected based on its census population size (*N*). The effective population size estimated for wildebeest in the GLE is 11% of the total estimated population size determined using aerial surveys in 2013 (*N* = 46,000) [35]—an effective to census population size ratio (*N*e/*N*) similar to estimates for many comparable mammals. Reviews have found the average *Ne*/*N* ratio in wildlife to be 0.10-0.14 [94,95]. In African buffalo, another large, highly mobile ungulate with a promiscuous mating system, researchers have estimated *N*e/*N* ratios ranging from 0.10 to 0.20 [96]. When also considering the moderate level of heterozygosity and low inbreeding coefficient for the Liuwa population, it seems unlikely that this estimate of current effective population size is indicative of a loss in genetic diversity after the population decline in the early 2000’s. Very recent population declines can actually increase the *N*e/*N* ratio, because census population size decreases faster than the timescale required for *Ne* to be reduced by a loss of genetic diversity [37,38,95]. By comparison, LaCava et al. [89] estimated a very low *N*e/*N* ratio of 0.01-0.02 for pronghorn (*Antilocaptra americana*), a North American ungulate that, in addition to having a polygynous mating system, experienced precipitous declines over a century ago.

Using both model-based and model-independent methods, we detected signatures of population expansion followed by decline in the site frequency spectrum of GLE blue wildebeest. Both the highest-likelihood expansion-decline fastsimcoal2 model and the stairway plot inferred a threefold increase in population size during the Middle Pleistocene, followed by an ∼87% reduction in population size during the Late Pleistocene and early Holocene. Evidence of Pleistocene population expansions followed by Holocene declines has been found in a number of other African mammal species using genetic data, including African buffalo [23,24], hippopotamus [26,27], African elephant [25,97], and savannah baboon [98]. In a comparative study of ruminant whole genomes, including 26 African species, Chen et al. [54] found using the pairwise sequentially Markovian coalescent (PSMC) method that most of the African species shared a similar demographic history of population expansion followed by decline in the late Pleistocene. Similarly, Liu et al. [39] used PSMC to infer effective population size over time for the five blue wildebeest subspecies, and the demographic history inferred for the brindled wildebeest (which did not include any individuals from the GLE) is very similar to the results of our stairway plot analysis in both the timing of the expansion and decline and the effective populations sizes inferred.

Two primary explanations have been proposed to explain the population declines in African ungulates during the late Pleistocene and early Holocene: climatic change and anthropogenic impacts [99]. Ice core records and other paleontological data suggest that Africa experienced large fluctuations in precipitation during the late Pleistocene and early Holocene, with extensive periods of drought [28–30]. Such climatic changes could have a profound impact on wildebeest, as modern-day wildebeest populations have fluctuated in response to drought [19] and wildebeest migration is at least partially driven by changes in rainfall and forage availability [4]. However, others have argued that climatic changes in the late Pleistocene were no more extreme than those occurring earlier in the epoch, and thus do not alone explain population declines and extinctions [99]. Human population growth and the advent of pastoralism during the Neolithic revolution may have increased pressures on ungulate populations already stressed by drought, leading to the observed declines [23,54]. Past population declines in the face of climatic changes and intensified human activity reflect the modern-day pressures faced by blue wildebeest, as habitat destruction, overhunting, and climate change all threaten the species today.

Our results show that—despite recent declines in population size and the disruption of migration to Angola—the GLE blue wildebeest population does not seem to have experienced recent reductions in genetic diversity or increased inbreeding. The ecosystem restoration efforts implemented in the GLE likely allowed the wildebeest population to increase before the bottleneck became severe enough to result in detectable reductions in genetic diversity.

However, wildebeest continue to face threats of habitat destruction, obstruction of migration routes from fencing, and climate change, and continued genetic monitoring could detect whether these impacts are influencing genetic diversity in the GLE in the future. This is especially important given the evidence that blue wildebeest population size drastically decreased during past periods of climate change and anthropogenic pressure in the late Pleistocene and early Holocene.

Future sampling and genomic analysis of wildebeest from Kafue National Park in Zambia could help to elucidate patterns of connectivity or differentiation between wildebeest in the GLE and those in a neighboring population that is also of conservation significance. The Zambezi River, which separates these two protected areas and currently acts as a barrier, may have historically prevented gene flow, leading to genetic differentiation between these populations.

Analysis of four wildebeest genomes from Kafue by Liu et al. [39] suggested that the Kafue population was genetically distinct from more southern brindled wildebeest populations, but the relationship between the Kafue and GLE populations remains unknown. Understanding patterns of genetic differentiation between GLE blue wildebeest and those in Kafue National Park would have important implications for future management decisions such as translocations between these populations. Our study provides a baseline understanding of genetic diversity and long- term demographic history for Africa’s second largest migratory blue wildebeest population, which will assist the development of effective conservation plans for the species as wildebeest face threats of habitat destruction and overhunting.

## Acknowledgements

Our thanks to the Zambia Department of National Parks and Wildlife, African Parks Network, and the Barotse Royal Establishment for permission and collaboration in this research. We gratefully acknowledge the late Jeffrey Horn of the NMU Computer Science Department, who passed away in February 2024. Jeff provided the computing infrastructure and assistance that made the data analysis for this work possible. We appreciate and miss his enthusiasm for collaborating across disciplines. We also thank Grant Combs for computing assistance. We are very grateful to Michael Sorenson and Jeffrey DaCosta for guidance on the RAD-seq protocol and analysis pipeline. We also thank Kurt Galbreath for input and advice on the project and Catherine Sun for feedback on the manuscript.

## Ethics Statement

All animal handling procedures were performed following animal welfare standards and protocols required by the Zambia Department of Veterinary and Livestock Services and the Zambia Department of National Parks and Wildlife and were approved by the Montana State University IACUC. Samples were exported with Zambia Wildlife Authority Permit #7830 and imported with USDA Veterinary Permit #109174.

## Author Contributions

KCT, ARL, MSB, and SJS contributed to the study conception and design. JM and ED collected the blue wildebeest tissues. HN and CC performed project support, oversight, and administration. RAD-seq library preparation and data analysis were performed by SJS. The first draft of the manuscript was written by SJS and all authors commented on the manuscript, contributed to subsequent drafts, and approved the final draft.

## Data Availability

The sequence data generated for this study are available from the NCBI Sequence Read Archive (SRA) under the following accession numbers: BioProject ID PRJNA588629; BioSample IDs SAMN13254976−SAMN13255074 (https://www.ncbi.nlm.nih.gov/bioproject/PRJNA588629).

