## Supplemental Materials for "Genetic diversity and demographic history of the largest remaining migratory population of blue wildebeest (*Connochaetes taurinus taurinus*) in southern Africa"

*PLOS ONE*

Stephanie J. Szarmach[1,2], Katherine C. Teeter[1], Jassiel M'soka[3], Egil Dröge[4,5], Hellen Ndakala[6], Clive Chifunte[4,7], Matthew S. Becker[4,8], Alec R. Lindsay[1]

[1]Department of Biology, Northern Michigan University, Marquette, MI 49855, USA

[2]Department of Biology, Pennsylvania State University, State College, PA 16802, USA

[3]USAID Zambia, Lusaka, Zambia

[4]Zambian Carnivore Programme, Mfuwe, Eastern Province, Zambia

[5]Wildlife Conservation Research Unit, Department of Biology, The Recanati-Kaplan Centre, Oxford University, Oxford, United Kingdom

[6]Zambia Department of National Parks and Wildlife, Liuwa Plain

[7]Zambia Department of National Parks and Wildlife, Mumbwa, Zambia

[8]Department of Ecology, Montana State University, Bozeman, MT 59735, USA

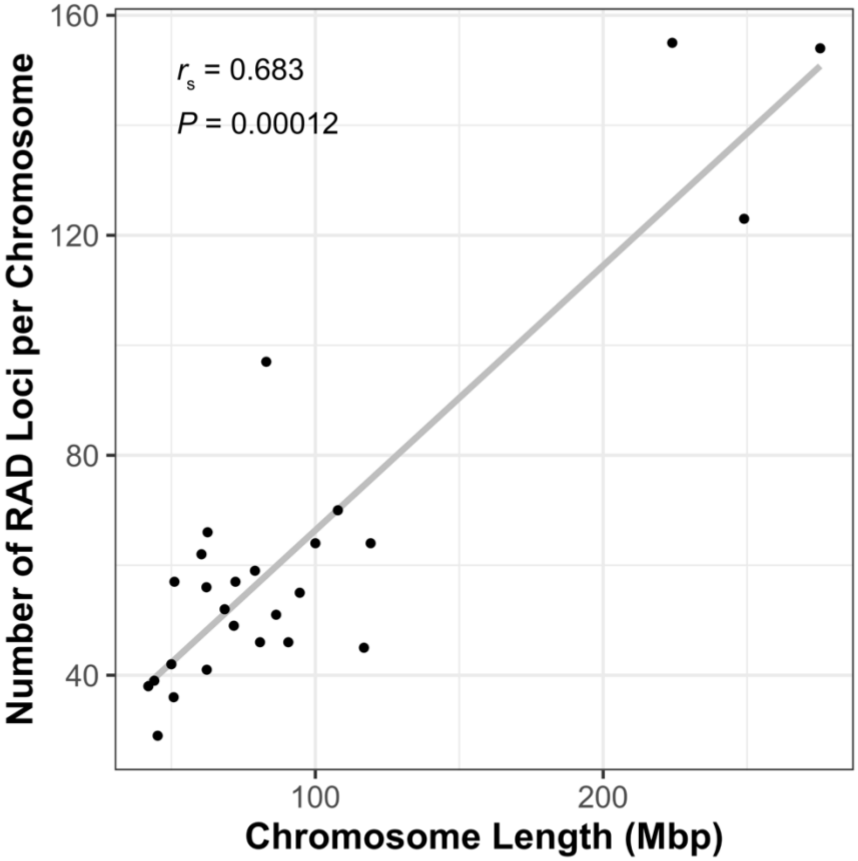


**Figure S1.** The number of RAD loci that aligned to each chromosome of the domestic sheep (*Ovis aries*) reference genome was significantly correlated with the length (Mbp) of the chromosome (Spearman's ρ = 0.683, $P$ = 0.00012). Linear regression line is shown in grey.

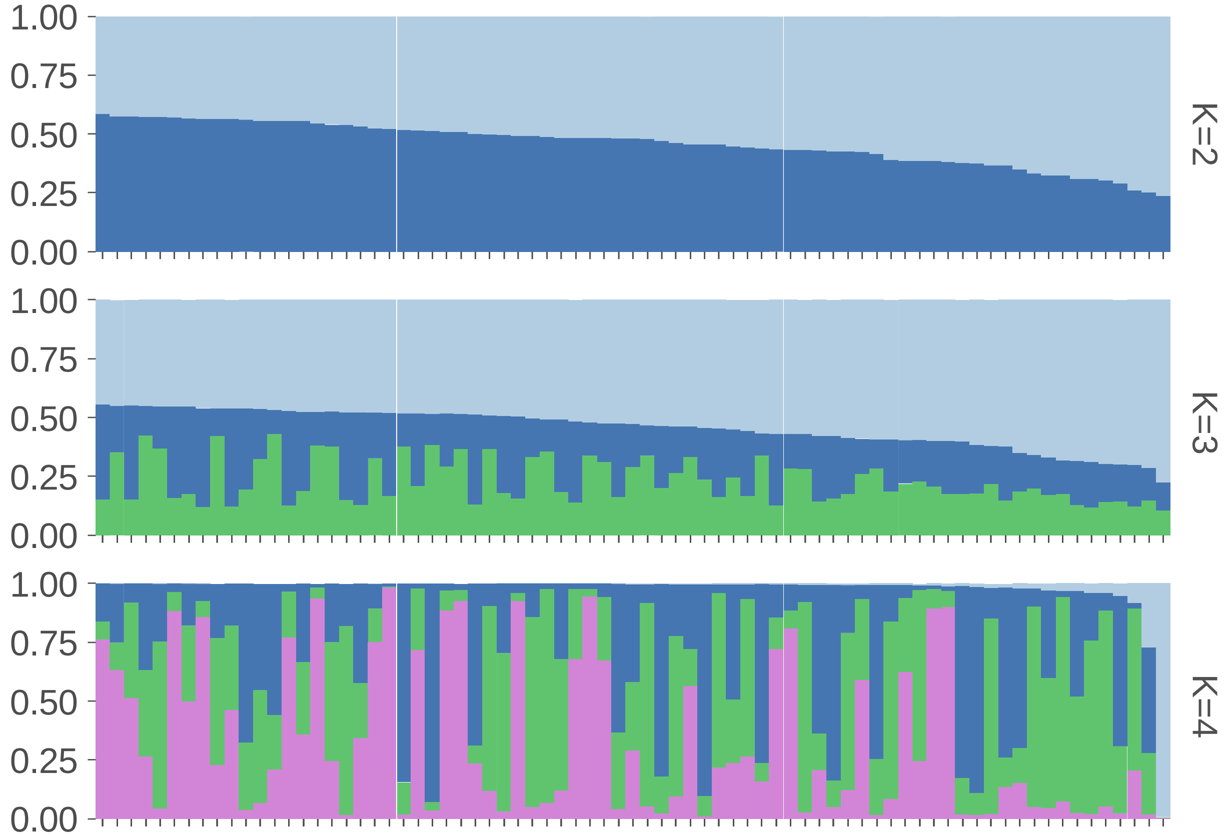


**Figure S2.** Structure plots for 75 wildebeest from the Greater Liuwa Ecosystem based on 1,730 SNPs assuming *K*=2, *K*=3, and *K*=4 clusters. None of these *K* values had a higher likelihood than *K*=1, and the assignment of most individuals to multiple clusters indicates that there is no genetic population structure within the GLE.

**Table S1.** Number of putative RAD loci remaining after each step of the quality filtering protocol. Data is presented separately for loci aligned to the domestic sheep (*Ovis aries*) or blue wildebeest (*Connochaetes taurinus*) reference genomes. Loci aligned to the wildebeest genome were used for downstream analyses.

| Step | Filtering requirement | Aligned to sheep reference genome | Aligned to wildebeest reference genome |
|---|---|---|---|
| 1 | Total putative loci | 19,437 | 19,459 |
| 2 | Contains ≥ 1 SNP | 12,404 | 12,430 |
| 3 | Genotyped in ≥ 90% of samples | 3,413 | 3,415 |
| 4 | < 5% genotypes with three alleles or a bad het. ratio | 2,986 | 2,986 |
| 5 | No loci with significant heterozygote excess | 2,967 | 2,965 |
| 6 | No high read depth outliers | 2,966 | 2,964 |
| 7 | One high-quality BLAST hit to reference genome[a] | 2,190 | 2,275 |
| 8 | No loci with BLAST hit to X-chromosome | 2,159 | 2,244 |
| 9 | No loci with significant homozygote excess | 2,096 | 2,160 |
| 10 | Meets expectations of infinite sites model | 2,017 | 2,027 |
| 11 | Problem loci identified through manual checks removed | 1,956 | 1,964 |
| 12 | Loci with ≤ 4 SNPs | 1,921 | 1,879 |
| 13 | Loci with no singleton SNPs (MAF > 0.01) | **1,773** | **1,730** |

[a]23 loci produced no hits to either the sheep or wildebeest genomes, but blastn searches resulted in hits to the genomes of related bovid taxa.

**Table S2.** Statistics obtained from Structure Harvester used to determine the optimal number of genetic clusters for wildebeest within the Greater Liuwa Ecosystem. For each *K* value (number of genetic clusters assumed in the model), we report the number of replicates, mean and standard deviation log probability of the data given the number of clusters, and Evanno statistic, Δ*K*.

| ***K*** | **Reps** | **Mean LnP(*K*)** | **SD LnP(*K*)** | **Δ*K*** |
|---|---|---|---|---|
| 1 | 10 | -87701.38 | 3.19 | NA |
| 2 | 10 | -94607.62 | 4652.74 | 1.28 |
| 3 | 10 | -95549.65 | 13257.36 | 0.51 |
| 4 | 10 | -89766.43 | 1509.91 | NA |

**Table S3.** Parameter estimates from the highest likelihood model for each demographic scenario tested in fastsimcoal2. Parameters describe effective population size (*N*) or the time a demographic event occurred (*T*) in years before present. Estimates have been rounded to the nearest hundred. The model with the highest support is shown in bold.

| **Model** | $N_{current}$ | $N_{ancestral}$ | $N_{intermediate}$ | $T_{expansion}$ | $T_{decline}$ |
|---|---|---|---|---|---|
| Constant size | 45,200 | - | - | - | - |
| Expansion | 44,100 | 41,900 | - | 1,501,300 | - |
| Decline | 9,800 | 53,400 | - | - | 4,000 |
| Bottleneck | 5,300 | 52,900 | 4,400 | - | 1,000 |
| **Expansion-Decline** | **13,400** | **26,800** | **92,700** | **635,600** | **15,480** |
